# Light quality signals generated by vegetation shade facilitate acclimation to reduced light quantity in shade-avoider plants

**DOI:** 10.1101/2020.09.14.296632

**Authors:** Luca Morelli, Sandi Paulišić, Wen-Ting Qin, Irma Roig-Villanova, Manuel Rodriguez-Concepcion, Jaime F. Martinez-Garcia

## Abstract

- When growing in search for light, plants can experience continuous or occasional shading by other plants. Plant proximity causes a decrease in the ratio of red to far red light (low R:FR) due to the preferential absorbance of red light and reflection of far red light by photosynthetic tissues of neighboring plants. This signal is often perceived before actual shading causes a reduction in photosynthetically active radiation (low PAR).
- Here we investigated elongation, photosynthesis and photoacclimation responses in several Brassicaceae species to explore the possible connections between low R:FR and low PAR.
- A negative correlation was found in shade-tolerant *Cardamine hirsuta* and shade-avoider *Arabidopsis thaliana* seedlings (e.g., shade-tolerance was associated with a good adaptation to low PAR but a poor or null response to low R:FR exposure). However, they could be genetically uncoupled. Most interestingly, exposure to low R:FR of shade-avoider plants improved their photoacclimation to low PAR by triggering changes in photosynthesis-related gene expression, pigment accumulation and chloroplast ultrastructure.
- These results indicate that low R:FR signaling unleashes molecular, metabolic and developmental responses that allow shade-avoider plants (including most crops) to adjust their photosynthetic capacity in anticipation of eventual shading by nearby plants.

## INTRODUCTION

Light is essential for plants as a source of energy and environmental information. Shading by nearby individuals can reduce light quantity (i.e. photon supply) and hence compromise photosynthetic activity and growth. To deal with this challenge, plants have developed response mechanisms based on the perception of light quality, i.e. spectral information (Martinez-Garcia *et al*., 2010; Casal, 2013). The preferential absorbance of red light (R) and reflection of far-red light (FR) by photosynthetic tissues results in a decreased ratio of R to FR (R:FR) when light is reflected from or filtered through green stems and leaves. The low R:FR is a very reliable light signal that announces the close presence of nearby plants that may compete for resources.

Plants growing in ecosystems where access to light is restricted (e.g. in forest understories) show a shade-tolerant habit by adapting their light capture and utilization systems to low light intensity conditions. By contrast, plants growing in open habitats are shade-avoiders (also referred to as shade-intolerant or sun-loving). In shade-avoider plant species, such as *Arabidopsis thaliana* and most sun-loving crops, perception of the low R:FR signal by the phytochrome photoreceptors activates a signaling pathway that eventually triggers a set of responses known as the shade avoidance syndrome (SAS). The most prominent phenotype following exposure to low R:FR is elongation (e.g., of seedling hypocotyl, leaf petiole and stem internode tissues), intended to overgrow neighboring competitors and outcompete them in the access to light. If the neighboring individuals overgrow and eventually shade the plant, the consequent reduction in light quantity (i.e., in the amount of radiation available for photosynthesis) results in additional and stronger SAS responses such as reduced leaf size, attenuated defense mechanisms and early flowering (Roig-Villanova & Martinez-Garcia, 2016).

The most extensively studied SAS response by far is hypocotyl elongation in *A. thaliana*. In this species, low R:FR inactivates phyB, releasing PHYTOCHROME INTERACTING FACTORs (PIFs) that can then regulate gene expression and promote elongation growth. This response is also repressed by negative SAS regulators such as ELONGATED HYPOCOTYL 5 (HY5), amongst many others (Cifuentes-Esquivel *et al*., 2013; Ciolfi *et al*., 2013). Biological activity of these transcription factors can be modulated by additional components of the SAS regulatory network such as LONG HYPOCOTYL IN FAR-RED 1 (HFR1, which binds PIFs to prevent their binding to target genes) and PHYTOCHROME A (phyA, which gets stabilized in shade and then promotes HY5 accumulation) (Ciolfi *et al*., 2013; Martinez-Garcia *et al*., 2014; Yang *et al*., 2018). Both HFR1 and phyA hence act as additional SAS repressors that were recently found to be instrumental for the adaptation to shade. Indeed, the shade-tolerant *Cardamine hirsuta*, a close relative of *A. thaliana*, does not elongate when exposed to low R:FR unless the function of phyA or HFR1 is genetically lost in mutant plants (Hay *et al*., 2014; Molina-Contreras *et al*., 2019; Paulisic *et al*., 2020).

Differences between shade-avoider and shade-tolerant species are not restricted to changes in elongation after exposure to low R:FR. *A. thaliana* and *C. hirsuta* also show a differential response to low R:FR in terms of photosynthetic pigment accumulation. Chlorophyll and carotenoid levels drop about 20 % in *A. thaliana* plants grown under low R:FR conditions, whereas the decrease is attenuated in *C. hirsuta* plants (Molina-Contreras *et al*., 2019). Photoacclimation (i.e., the ability of plants to respond with specific phenotypic adjustments to changes in the incident light) also diverges. The shade-avoider *A. thaliana* showed a lower capacity to acclimate to reduced photosynthetically active radiation (low PAR) but a higher capacity to acclimate to intense light (high PAR) compared to the shade-tolerant *C. hirsuta* (Molina-Contreras *et al*., 2019). However, the possible connections between low R:FR signaling and photoacclimation responses remain virtually unknown. Here we explored natural and engineered genetic diversity to investigate this connection.

## MATERIALS AND METHODS

### Plant material and growth conditions

*Arabis alpina* (*pep1-1* mutant) (Wang *et al*., 2009), *Arabidopsis thaliana* (Col-0 accession), *Cardamine hirsuta* (Oxford, Ox accession) (Molina-Contreras *et al*., 2019), *Capsella bursa-pastoris* (accessions Strasbourg-1, Str-1 and Freiburg-1, Fre-1), *Capsella rubella* and *Sisymbrium irio* plants were grown in the greenhouse under long-day photoperiods (16 h light and 8 h dark) to produce seeds, as described (Gallemi *et al*., 2017). Seeds of *C. bursa-pastoris* were collected by Ruben Alcazar (University of Barcelona, Spain) from wild populations in Strasbourg (France, coordinates: 48.612436, 7.767881; Str-1) and Freiburg (Germany, coordinates: 47.994945, 7.861979; Fre-1). Seeds of *Capsella rubella*, collected from wild populations in Crete (Greece, coordinates 35.29, 24.42; accession 879) were previously described (Koenig *et al*., 2019). Seeds of *Sisymbrium irio* were collected from wild populations in Bellaterra (Barcelona, Spain, coordinates: 41.497731, 2.109558). Seeds of *Nasturtium officinale* were provided by a seed company (www.semillasfito.es). *A. thaliana* and *C. hirsuta* mutant and transgenic lines were previously available in our laboratories (Molina-Contreras *et al*., 2019; Ortiz-Alcaide *et al*., 2019; Paulisic *et al*., 2020).

For the light acclimation experiments seedlings were germinated and grown on medium without sucrose: 2.2 g/L MS basal salt mixture (Duchefa), 1% (w/v) agar, 0.25 g/L 2-(*N*-morpholino)ethanesulfonic acid -MES-(Sigma Aldrich), pH 5.7). Normal light conditions (W) refer to light produced by cool-white vertical fluorescent tubes (PAR of 20-24 µmol m^-2^ s^-1^) with a R:FR of 1.5-3.3. Low light (L) and high light (H) conditions corresponded to PAR values of 4 and 200 µmol m^-2^ s^-1^, respectively produced by horizontal fluorescent tubes. Low R:FR treatment was produced by supplementing W with FR (W+FR). FR was emitted from a GreenPower LED module HF far-red (Philips), providing a R:FR of 0.02 (Martinez-Garcia *et al*., 2014). Light fluence rates were measured with a Spectrosense2 meter (Skye Instruments Ltd); full spectra photon distribution of W and W+FR treatments have been described elsewhere (Molina-Contreras *et al*., 2019).

### Measurement of hypocotyl length

For hypocotyl measurement, about 30 seeds of each genotype were germinated and grown on plates. For quantification of hypocotyl length, at least 20 seedlings were analyzed with the FIJI-ImageJ software (Schindelin *et al*., 2012), as described (Roig-Villanova *et al*., 2019). All experiments were repeated at least three times with consistent results. Hypocotyl measurements from all the different experiments were averaged.

### Photosynthetic measurements and pigment quantification

Whole seedlings were harvested, ground in liquid nitrogen, and the resulting powder was used for quantification of chlorophylls and carotenoids either spectrophotometrically or by HPLC as described (Bou-Torrent *et al*., 2015). Chlorophyll fluorescence measurements were carried out on seedlings using a MAXI-PAM fluorometer (Heinz Walz GmbH) as described (Molina-Contreras *et al*., 2019). Briefly, for every measurement the whole cotyledons of 7 seedlings were considered. Effective quantum yield of photosystem II (PSII) under growth light, ϕPSII, was measured as ΔF/Fm’, where ΔF corresponds to Fm’-F (the maximum minus the minimum fluorescence of light-exposed plants). Maximum quantum yield of PSII, Fv/Fm, was calculated as (Fm-Fo)/Fm, where Fm and Fo are respectively the maximum and the minimum fluorescence of dark-adapted samples. For dark acclimation, plates were incubated for at least 30 minutes in darkness to allow the full relaxation of photosystems. Light curves were constructed with 10 incremental steps of actinic irradiance (E; 0, 20, 55, 110, 185, 280, 395, 530, 610, 700 μmol photons·m^-2^·s^-1^of PAR). For each step, ϕPSII was monitored every minute and relative electron transport rate (rETR) was calculated as E×ϕPSII. The light response and associated parameters ETRm (maximum electron transport rate) and alpha (photosynthetic rate in light-limited region of the light curve) were characterized by fitting iteratively the model of the rETR versus E curves using MS Excel Solver (Platt *et al*., 1980). The fit was very good in all the cases (r>0.98).

### Microarray data analyses

Microarray data corresponding to *A. thaliana* Col-0 (At-WT) and At-*pifq* seedlings exposed to low-R:FR for 0, 1, 3 and 24 h (Leivar *et al*., 2012) were analyzed to select for differentially-expressed genes (DEGs) specifically related to photosynthesis. The reported list of DEGs was further filtered using cut-offs of FDR <0.05 and log2-transformed fold change (log2FC) higher than 0.585 for upregulated genes and lower than -0.599 for downregulated genes. Then, photosynthesis-related genes were identified by using the KEGG (Kyoto Encyclopedia of Genes and Genomes) Mapper tool (Kanehisa & Sato, 2020).

### Transmission electron microscopy

Transmission electron microscopy (TEM) was carried out as described (Flores-Perez *et al*., 2008). Chloroplast features in the pictures were quantified by using the FIJI-ImageJ software (Schindelin *et al*., 2012).

## RESULTS

### Different Brassicaceae species present divergent photoacclimation responses

We previously showed that, compared to sun-loving *A. thaliana* Col-0 (At), shade-tolerant *C. hirsuta* Ox (Ch) exhibits a better ability to maintain photosynthesis after transfer to low PAR but a stronger chlorophyll loss when light intensity increases (Molina-Contreras *et al*., 2019). To better characterize the photoacclimation responses of these two Brassicaceae species, both At and Ch were germinated and grown for 7 days under control PAR conditions (W, 20-24 µmol m^-2^ s^-1^) and then transferred to either low PAR (L, 4 µmol m^-2^ s^-1^) or high PAR (H, 200 µmol m^-2^ s^-1^) for 7 more days (Fig. 1). Light curve analysis at day 3 after the transfer showed a better photosynthetic activity of Ch compared to At when grown under L and a better activity of At compared to Ch when grown under H (Fig. 1a). Derived parameters such as ETRm (maximum electron transport rate) and alpha (alpha, photosynthetic rate in light-limited region of the light curve) also illustrated that At performed better than Ch after transfer to H but worst after transfer to L (Fig. 1b). Other photosynthetic parameters such as maximum quantum efficiency of PSII (Fv/Fm) and light use efficiency of PSII (ϕPSII) only showed differences between At and Ch at longer times of exposure to either H or L (Fig. 1c). At day 7, Fv/Fm values were lower in Ch than in At after transfer to H, while the opposite was observed when transferred to L. A similar trend was observed in the case of ϕPSII (Fig. 1c). These results together indicate that Ch tolerates better the transfer to low PAR (consistent with Ch being more tolerant to shade), while an increase in light irradiance compromises photosynthetic efficiency in Ch more than in shade-avoider At. Based on these results, we used light curve analysis to estimate photoacclimation to low PAR and Fv/Fm measurements to estimate photoacclimation to high PAR.

**Figure 1.**
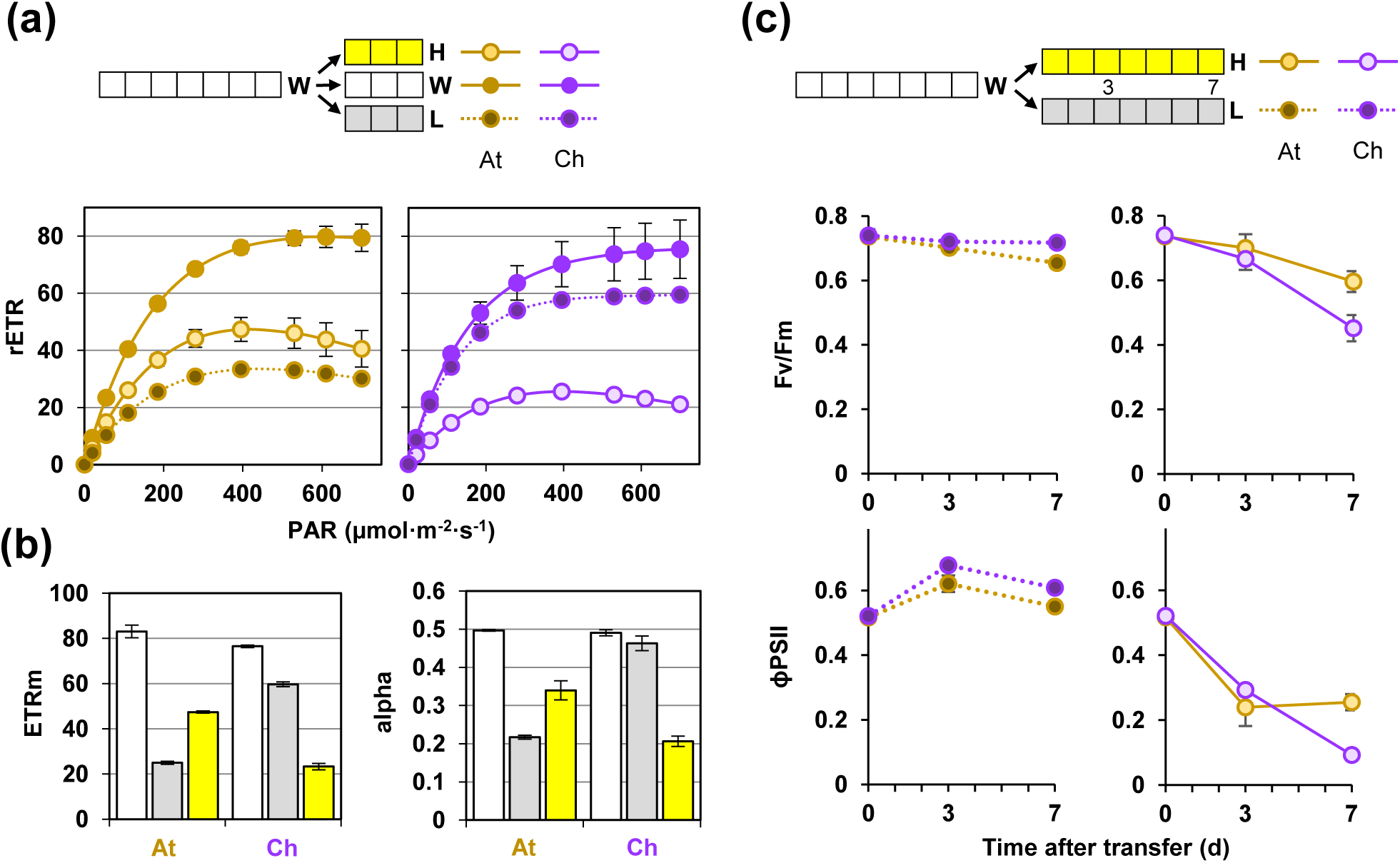
*Arabidopsis thaliana* and *Cardamine hirsuta* show antagonistic photoacclimation responses to high and low PAR. **(a)** Light curves of *A. thaliana* (At) and *C. hirsuta* (Ch) seedlings germinated and grown under normal white light (W) for 7 days and then either kept under W or transferred to either high light (H) or low light (L) for 3 more days. Values represent the mean and standard error of n=3 plants for treatment. **(b)** Maximum relative electron transport rate (ETRm) and photosynthetic rate in the light-limited region of the light curve (alpha) calculated from the curves shown in (a). **(c)** Maximum photochemical efficiency of PSII in the dark-adapted state (Fv/Fm) and effective quantum yield calculated at growth light (ϕPSII) of seedlings germinated and grown for 7 days under W and then transferred to either H or L for more 7 days. Data were taken at 0, 3 and 7 days after the transfer.Values are mean and standard error of n=9 seedlings per treatment.

Besides At and Ch, the Brassicaceae family (mustards) includes many food crops (e.g., cauliflower, broccoli, radish, cabbage, kale, and similar green leafy vegetables) and a diversity of wild species from forested and open habitats. As a first step to explore the possible connection between low PAR and low R:FR responses, we analyzed photoacclimation and hypocotyl elongation in six different Brassicaceae species or accessions, including At and Ch as controls. The selected wild mustards were *Arabis alpina* (Aa), two accessions of *Capsella bursa-pastoris*, Freiburg-1 (Cb-F) and Strasbourg-1 (Cb-S), *Capsella rubella* (Cr), *Nasturtium officinale* (No), and *Sisymbrium irio* (Si). Initially, we aimed to classify them as shade-avoider or shade-tolerant based on photoacclimation responses. After germination and growth for 7 days under W, seedlings were either kept under W or transferred to L for one additional day before performing light curve analyses (Fig. 2). Similar to the shade-avoider At, seedlings of Cb-F, Cb-S and Cr showed a lowering of the curve under L conditions, whereas those of Aa, No and Si behaved as the shade-tolerant Ch and showed virtually identical light curves under W and L (Fig. 2a). ETRm and alpha values also illustrated that the L treatment led to decreased photosynthetic performance in At, Cb-F, Cb-S and Cr but not in Ch, Aa, No and Si (Fig. 2b). We next analyzed photoacclimation to increased irradiation quantifying Fv/Fm before or after transferring 7-day-old W-grown seedlings to H for 7 additional days. Again, At grouped together with the two accessions of Cb and with Cr as they acclimated much better to high PAR compared to the group formed by Ch, Aa, No and Si (Fig. 2c). Together, these photoacclimation results led to classify the former group as shade-avoiders, and the latter as shade-tolerant species.

**Figure 2.**
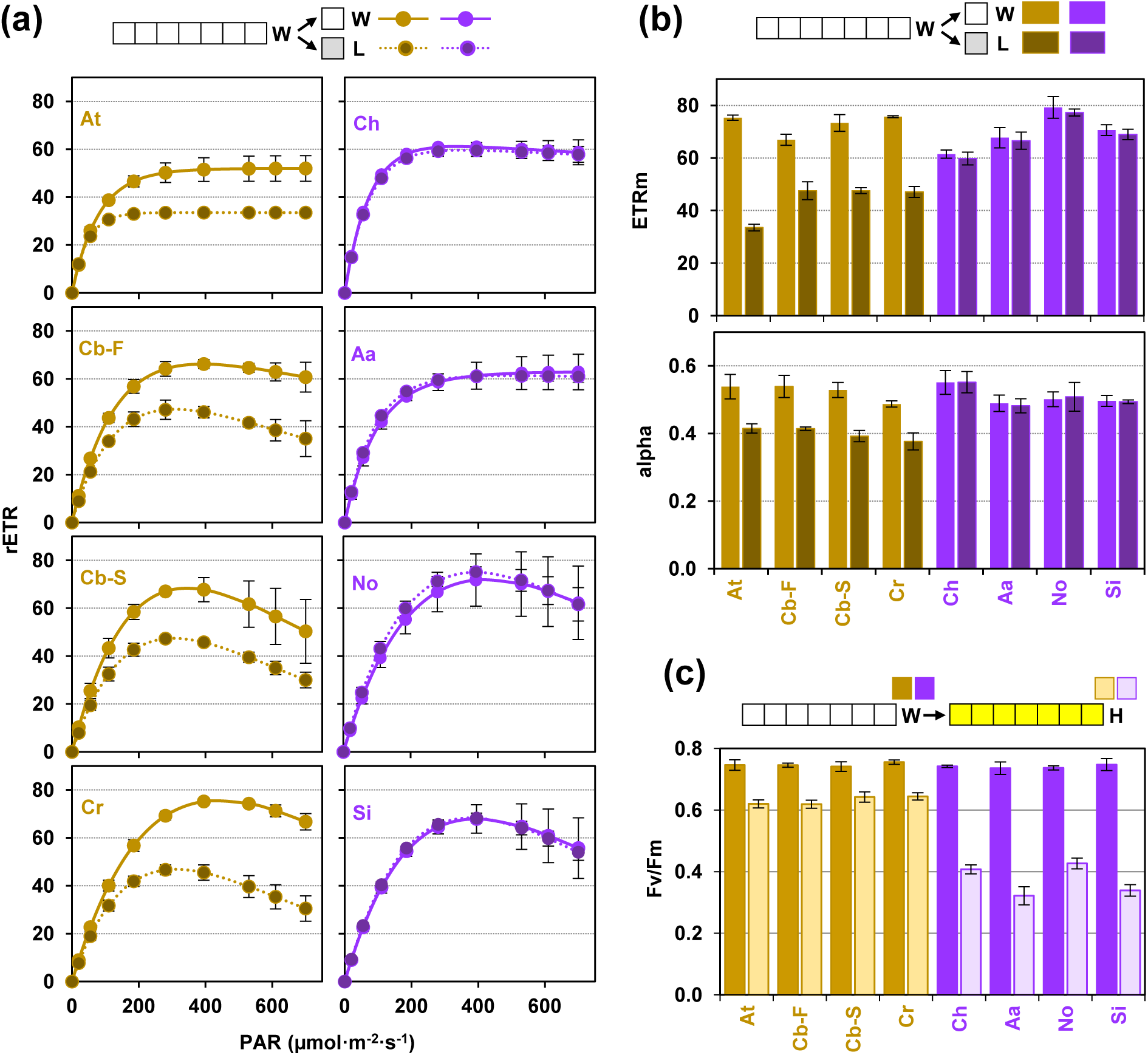
Brassicaceae plants can be grouped with either *Arabidopsis thaliana* or *Cardamine hirsuta* based on their photoacclimation responses. **(a)** Light curves of *Arabidopsis thaliana* (At), *Capsella bursa-pastoris* (Cb-F and Cb-S), *Capsella rubella* (Cr), *Cardamine hirsuta* (Ch), *Arabis alpina* (Aa), *Nasturtium officinale* (No), and *Sysimbrium imbrio* (Si) seedlings germinated and grown under normal white light (W) for 7 days and then either kept under W or transferred to low light (L) for 1 more day. Values represent the mean and standard error of n=3 plants for treatment. **(b)** ETRm and alpha values calculated from the curves shown in (a). **(c)** Fv/Fm values of seedlings grown for 7 days under W and then transferred to high light (H) for 7 more days. Mean and standard error of n=9 seedlings per treatment are represented.

### Photoacclimation responses can be uncoupled from shade-driven hypocotyl elongation

Next, we investigated whether the classification of the selected mustard species as shade-avoider or shade-tolerant based on their photoacclimation features correlated with their elongation response to low R: FR. After germination and growth for 3 days under W (R:FR=1.5-3.3), seedlings were either kept under W or transferred to FR-supplemented W (W+FR, R:FR=0.02) for 4 additional days, and then hypocotyl length was measured (Fig. 3). Similar to At, the Cb-F accession showed a strong hypocotyl elongation response, whereas Cb-S, Cr and No elongated moderately in response to low R: FR. By contrast, Ch, Aa and Si did not elongate in response to low R:FR (Fig. 3a). These results indicate that the elongation response to low R:FR cannot be fully predicted based on the photoacclimation phenotype of a particular accession. Nonetheless, accessions classified as shade-avoider based on their photoacclimation behavior (i.e. poor photoacclimation to decreased PAR but good photoacclimation to increased PAR) exhibit a range of responses to low R:FR (i.e. from moderate to strong elongation), whereas plant species with a shade-tolerant photoacclimation responses display either no elongation or a mild shade-avoider phenotype in terms of hypocotyl elongation when exposed to low R:FR (e.g. No). These data together suggest that low R:FR-induced hypocotyl elongation and PAR photoacclimation responses do not fully correlate in nature.

**Figure 3.**
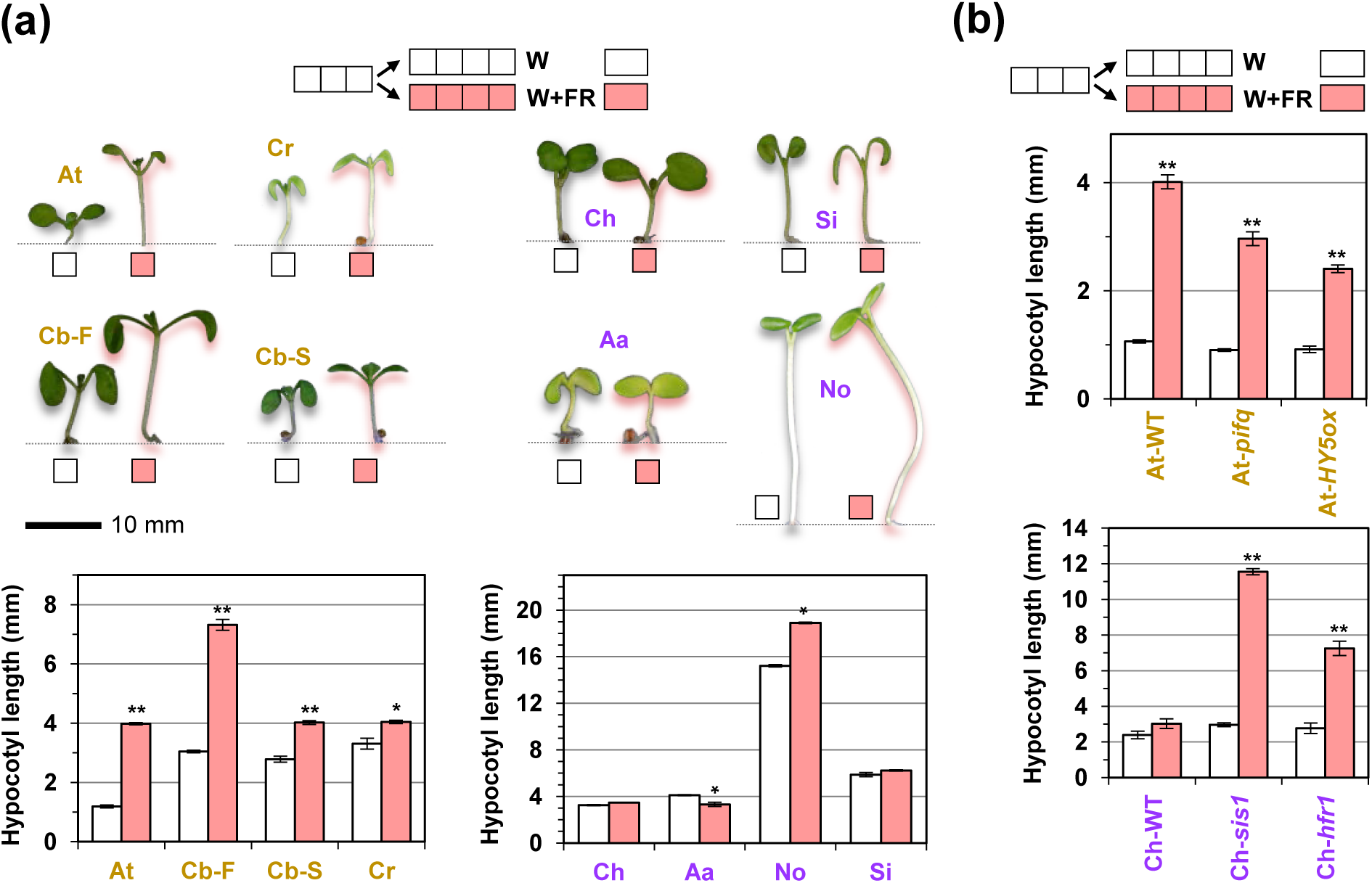
The hypocotyl elongation response to low R:FR is plastic in Brassicaceae plants. **(a)** The indicated genotypes were germinated and grown under normal white light (W) for 3 days and then either kept under W or transferred to low R:FR (W+FR) for 4 more days. Then, pictures were taken and hypocotyl length was measured. **(b)** Hypocotyl length of the indicated mutants grown as indicated in (a). In both (a) and (b), mean and standard error of measurements from at least 20 seedlings in n=3 independent experiments per treatment are represented. Asterisks mark statistically significant changes in W+FR relative to W (*t* test, * *P*<0.05 and ** *P*<0.01).

The shade-avoider or shade-tolerant elongation phenotype in response to low R:FR can be reversed by manipulating the levels of specific SAS regulators. Previous results have shown that At lines overexpressing *HY5* (At-*HY5ox*) display an attenuated hypocotyl response to low R:FR (Ortiz-Alcaide *et al*., 2019), whereas a similar but weaker response was observed in a quadruple mutant defective in all members of the photolabile PIF quartet (At-*pifq*) (Fig. 3b). Despite the different degrees of elongation response to low R:FR, these two lines showed photoacclimation responses to low PAR very similar to those of wild-type (At-WT) controls (Fig. 4). Both light curves (Fig. 4a) and ETRm values (Fig. 4b) were almost identical in At-WT plants and mutants hyposensitive to low R:FR (Fig. 3b). In the case of *C. hirsuta*, lines deficient in phyA (Ch-*sis1*) or HFR1 (Ch-*hfr1*) gain the ability to elongate when exposed to low R:FR (Molina-Contreras *et al*., 2019; Paulisic *et al*., 2020) (Fig. 3b). In contrast to the shade-hyposensitive At mutants, the hypersensitive Ch mutant lines appeared to gain a partial shade-avoider phenotype in terms of photoacclimation to low PAR, as a trend towards lower values of light curves (Fig. 4a) and ETRm (Fig. 4b) were observed under L compared to W. However, photoacclimation to increased PAR estimated from Fv/Fm values and also from chlorophyll levels (Molina-Contreras *et al*., 2019) was similar for Ch-WT, Ch-*sis1* and Ch-*hfr1* plants (Fig. 4c). We therefore concluded that manipulation of the plant ability to elongate in response to proximity shade hardly impacts their photoacclimation capacity, at least when plants are growing in the absence of the low R:FR signal.

**Figure 4.**
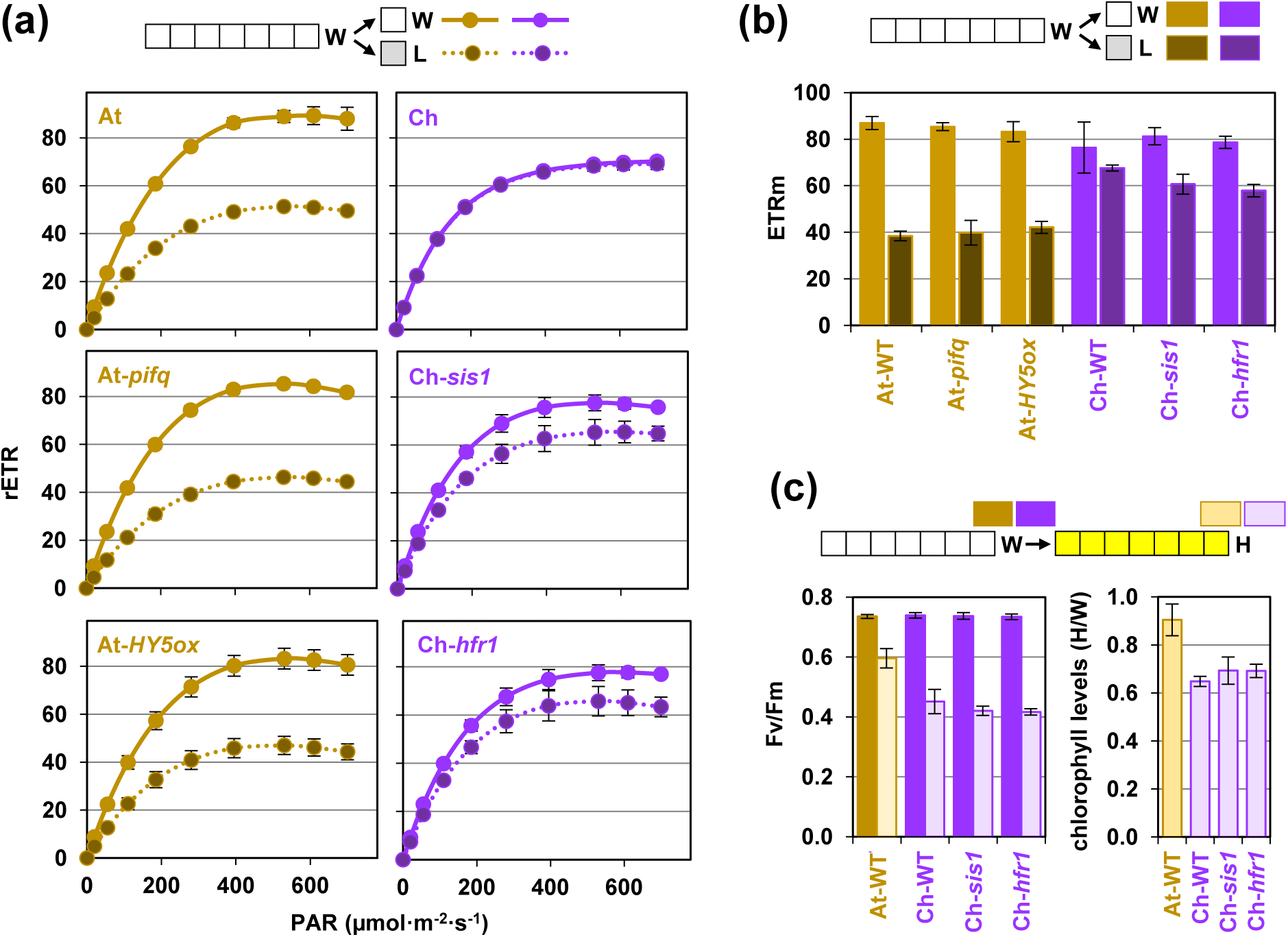
Mutations that alter sensitivity to low R:FR do not impact photoacclimation responses. **(a)** Light curves of *A. thaliana* and *C. hirsuta* wild-type and mutant seedlings germinated and grown under normal white light (W) for 7 days and then either kept under W or transferred to low light (L) for 1 more day. Values represent the mean and standard error of n=3 plants for treatment. **(b)** ETRm values calculated from the curves shown in (a). **(c)** Fv/Fm values and HPLC-determined chlorophyll levels of seedlings grown for 7 days under W and then transferred to high light (H) for 7 more days. Mean and standard error of n=9 seedlings (Fv/Fm) or n=3 independent pools (HPLC) per treatment are represented.

### Activation of low R:FR signaling causes a decrease in pigment levels and photosynthetic activity

Low R:FR signals not only influence hypocotyl elongation but they are also known to reduce the contents of photosynthetic pigments (chlorophylls and carotenoids) in many plant species (Roig-Villanova *et al*., 2007; Cagnola *et al*., 2012; Patel *et al*., 2013; Bou-Torrent *et al*., 2015; Molina-Contreras *et al*., 2019). The reduction is observed in both elongating (At-WT) and non-elongating (Ch-WT) seedlings, but it is stronger in the former (Fig. 5). *C. hirsuta* mutants that gained the ability to elongate in response to shade, such as Ch-*sis1* and Ch-*hfr1*, also displayed stronger reductions in photosynthetic pigment contents relative to Ch-WT after low R:FR exposure (Fig. 5a) (Molina-Contreras *et al*., 2019). Conversely, *A. thaliana* mutants with a reduced ability to elongate in response to shade, such as At-*pifq* and At-*HY5ox* (Fig. 3b), showed attenuated reduction of pigment contents relative to At-WT when exposed to low R:FR (Fig. 5a).

**Figure 5.**
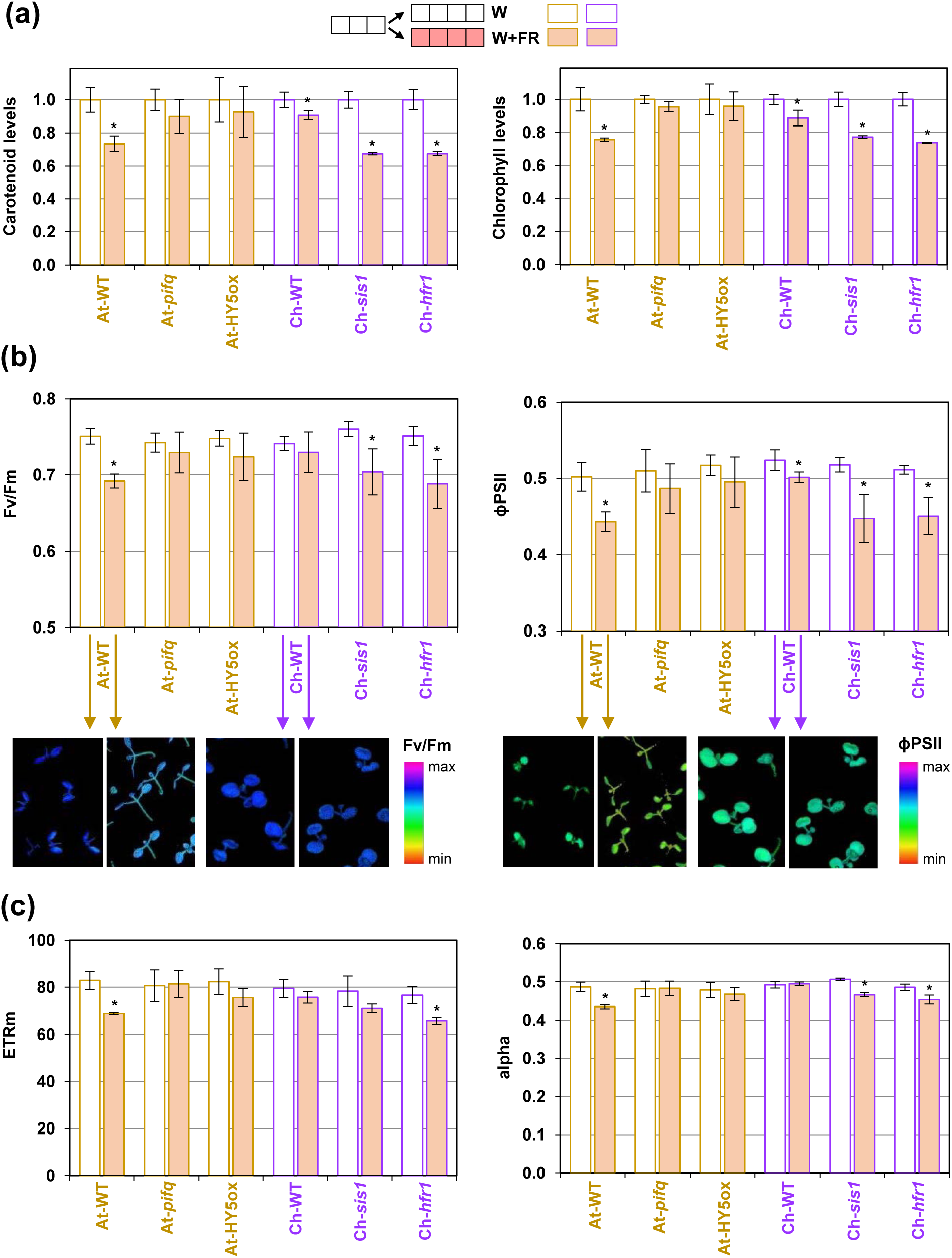
Activation of low R:FR signaling reduces photosynthetic pigment levels and activity. **(a)** The indicated genotypes were germinated and grown under normal white light (W) for 3 days and then either kept under W or transferred to low R:FR (W+FR) for 4 more days. Then, the levels of photosynthetic pigments (carotenoids and chlorophylls) were quantified spectrophotometrically. **(b)** Fv/Fm and ϕPSII values of seedlings germinated and grown as indicated in (a). Lower pictures show false-color images of the indicated parameters in wild-type seedlings. **(b)** ETRm and alpha values of seedlings germinated and grown as indicated in (a). In the plots, mean and standard error of n=3 independent pools of seedlings (a) or n=9 seedlings (b, c) per treatment are represented. Asterisks mark statistically significant changes in W+FR relative to W (*t* test, *P*<0.05).

To test whether decreases in photosynthetic pigment levels driven by simulated shade exposure might affect photosynthetic activity, we next measured Fv/Fm and ϕPSII in seedlings grown either under W or under W+FR (Fig. 5b). Indeed, low R:FR was found to result in decreased photosynthetic activity in the lines with strong pigment loss responses independently on the species (At-WT, Ch-*sis1* and Ch-*hfr1*). ETRm and alpha parameters also tended to be lower in W+FR-exposed At-WT, Ch-*sis1* and Ch-*hfr1* seedlings compared to W controls (Fig. 5c). The effect of low R:FR on photosynthesis was much less dramatic in the rest of the lines (At-*pifq*, At-*HY5ox* and Ch-WT), which consistently displayed a reduced impact of W+FR exposure on their photosynthetic pigment levels (Fig. 5).

Proximity shade signals have also been found to impact photosynthesis at the level of gene expression. Analyses of low R:FR-triggered transcriptomic changes showed reduced levels of transcripts encoding photosynthesis-related proteins (e.g. enzymes involved in chlorophyll and carotenoid biosynthesis, components of the photosynthetic apparatus, and/or members of the carbon fixation process) in several species, including alfalfa (Lorenzo *et al*., 2019), maize (Shi *et al*., 2019), tomato (Cagnola *et al*., 2012) and *A. thaliana* (Leivar *et al*., 2012). Interestingly, the changes in the expression of photosynthesis-related genes triggered by low R:FR are attenuated in the At-*pifq* mutant compared to At-WT seedlings (Fig. 6). This is particularly evident in the case of low R:FR-repressed photosynthetic genes (Fig. 6), suggesting that the PIF-mediated regulation of gene expression in response to low R:FR is instrumental for the observed changes in photosynthesis (Fig. 5).

**Figure 6.**
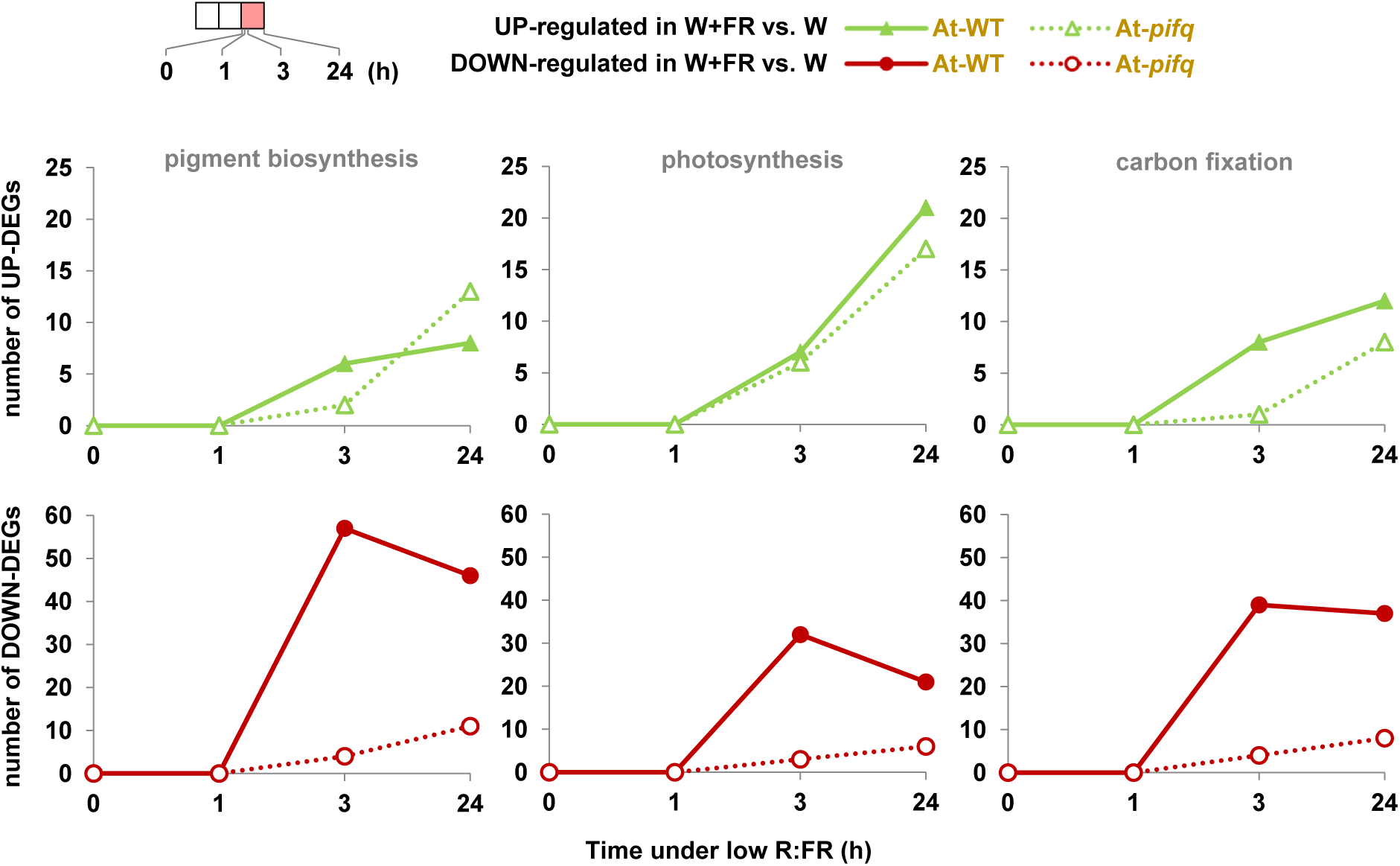
Exposure to low R/FR triggers changes in photosynthetic gene expression that are attenuated in the hyposensitive At-*pifq* mutant. Data were extracted from a publicly available experiment (Leivar *et al*., 2012). At-WT and At-*pifq* lines were germinated and grown under normal white light (W) for 2 days and exposed to low R:FR (W+FR) for 0, 1. 3 or 24 h. Plots represent the number of differentially expressed genes (DEGs) either up-regulated or down-regulated in W+FR vs. W that are involved in photosynthetic pigment biosynthesis (KEGG pathways ath00906 and ath00860), photosynthesis (ath00195 and ath00196), and carbon fixation (ath00710).

### Exposure of shade-avoider plants to low R:FR improves their photoacclimation to low PAR

The observation that exposure of low R:FR caused a decreased in photosynthetic activity of At-WT seedlings and shade-hypersensitive Ch mutants prompted us to analyze whether this light signal may also cause changes in chloroplast ultrastructure. Cotyledons from At-WT seedlings germinated and grown for 2 days under W and then either kept in W or transferred to W+FR for 5 additional days were collected and used for transmission electron microscopy (TEM). Chloroplasts from low R:FR-exposed samples were found to exhibit larger grana stacks and contain less and smaller plastoglobules compared to W-grown controls (Fig. 7). Interestingly, similar changes are associated to low PAR photoacclimation (Rozak *et al*., 2002; Lichtenthaler, 2007; Wood *et al*., 2018). We therefore reasoned that exposure to low R:FR in the absence of any light intensity change might trigger responses to anticipate a foreseeable shading involving a decrease in PAR. To test this hypothesis, we analyzed light curves of WT and mutant seedlings grown in either W or W+FR and then transferred to low PAR for 3 days (Fig. 8). Pre-exposure of At-WT seedlings to low R:FR (W+FR) resulted in a strongly attenuated reduction in ETRm and alpha after their transfer to low PAR (Fig. 8a). By contrast, At mutants with reduced SAS elongation responses also lost the response to low R:FR in terms of improved photoacclimation to L (Fig. 8a). Pre-treatment with W+FR had virtually no effect on the photoacclimation of Ch-WT seedlings to low PAR but caused a slight improvement of ETRm in shade-hypersensitive Ch mutants at day 1 after transfer to L (Fig. 8a). When analyzing photoacclimation to high PAR, pre-exposure of At-WT or Ch-WT seedlings to W+FR resulted in no improvement compared to W-grown controls (Fig. 8b). If anything, Ch-WT seedlings grown under W+FR photoacclimated worse than W-grown seedlings when exposed to higher light intensity (Fig. 8b). Based on these data we conclude that detection and transduction of low R:FR signals not only allows plants to overgrow their neighbors but also to pre-adapt their photosynthetic machinery to foreseeable conditions of actual shading involving reduced PAR. By contrast, shade-tolerant plants have a better adapted photosynthetic capacity to grow under reduce PAR and do not seem to use the low R:FR signal.

**Figure 7.**
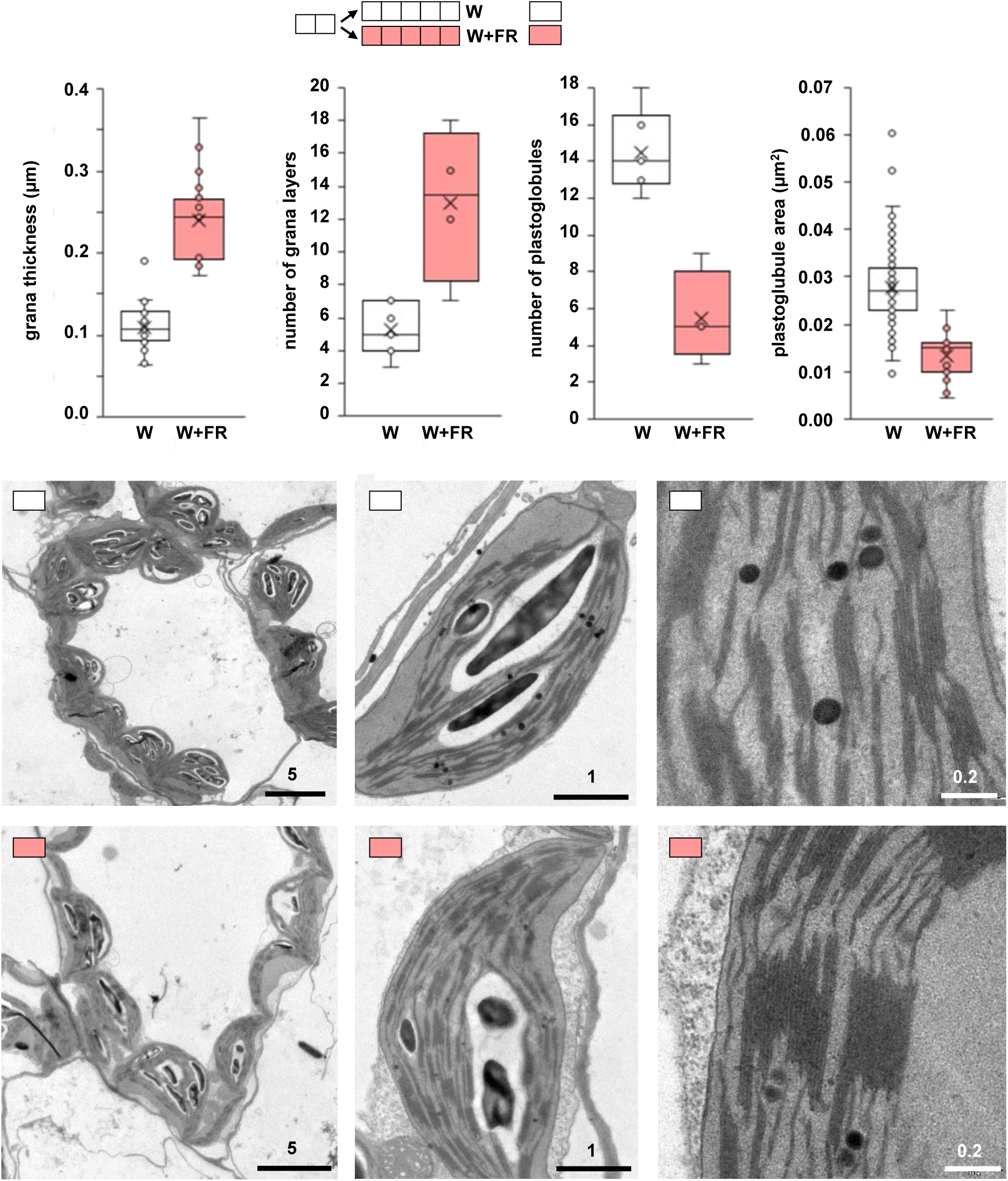
Low R:FR triggers ultrastructural changes in *A. thaliana* chloroplasts. At-WT seeds were germinated and grown under normal white light (W) for 2 days and then either kept under W or transferred to low R:FR (W+FR) for 5 more days. Cotyledons were then used for TEM analysis of chloroplast ultrastructure. Representative pictures at different scales (numbers indicate µm) are shown. Boxplots show quantification of the indicated parameters from the images.

**Figure 8:**
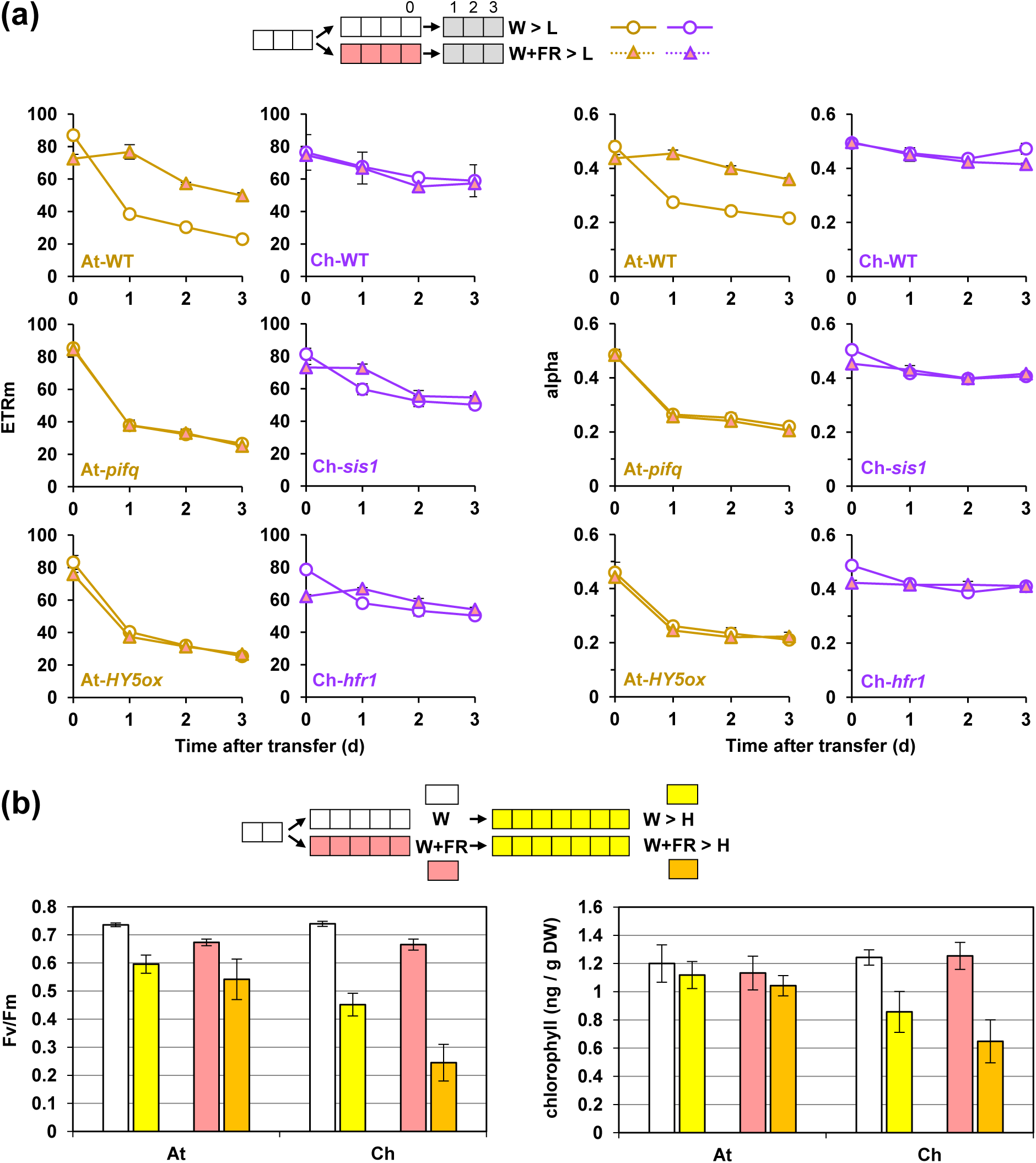
Pre-exposure to low R:FR improves the photoacclimation to low PAR in shade-avoider plants. **(a)** The indicated genotypes were germinated and grown under normal white light (W) for 3 days, transferred to either W or W+FR for 4 days, and then exposed to L for 3 more days. Mean and standard error of ETRm and alpha values at 0, 1, 2 and 3 days after transfer to L are shown (n=9 seedlings per treatment). **(b)** Wild-type *A. thaliana* and *C. hirsuta* lines were germinated and grown under W for 2 days, transferred to either W or W+FR for 5 days, and then exposed to H for 7 more days. Fv/Fm values and HPLC-determined chlorophyll levels were determined. Mean and standard error of n=9 seedlings (Fv/Fm) or n=3 independent pools (HPLC) per treatment are represented.

## DISCUSSION

Plants have been traditionally classified as shade avoider and tolerant based mostly on their natural habitat, although virtually all plants are exposed to at least some degree of shade during their lifetime. As an ecological concept, shade tolerance refers to the capacity of a given plant to tolerate low light levels, but it is also associated with a wide range of traits, including phenotypic plasticity to optimize light capture (Valladares & Niinemets, 2008). Analyzing a range of caulescent herbs, it was suggested that the elongation response upon exposure to low R:FR was dependent on the shade habit, the shade-avoiders elongating the most and the shade-tolerant showing a mild or no elongation response (Smith, 1982). Indeed, elongation might not be the best solution for plants that spend all their lives under a canopy or permanently shaded by other plants. Another important parameter to ascertain the degree of shade tolerance of a plant is photoacclimation capacity, which is essential for plant fitness in environments with changing light input conditions (e.g., those where the growth of nearby plants may suddenly compromise access to light). By taking into account both parameters (the hypocotyl elongation response and the capacity to acclimate to low or high PAR), here we analyzed the shade tolerance of several Brassicaceae species, including the closely related mustard model systems *A. thaliana* and *C. hirsuta*. As a rule of thumb, we observed that *C. hirsuta* and other species showing a good photoacclimation response to low PAR (and badly performing after transfer to high PAR) showed a poor or null elongation response to low R:FR (Fig. 2, 3).

Mustards such as *A. thaliana* that photoacclimated poorly to low PAR but better to high PAR tended to more conspicuously elongate their hypocotyls in response to low R:FR, but there were exceptions of poorly elongating species such as *Nasturtium officinale* (Fig. 2, 3). Furthermore, mutation of genes encoding SAS regulators can dramatically change the elongation response to low R:FR without improving the photoacclimation phenotype (Fig. 4). Besides confirming that the capacity for photosynthetic acclimation to changing irradiance is species-specific (Bailey *et al*., 2001), our results support the conclusion that measuring the hypocotyl elongation response to low R:FR is not sufficient *per se* to classify a species as shade-tolerant or shade-avoider. Instead, the photoacclimation response to low and high PAR appears to be a better indicator of shade tolerance. A combination of both the shade-induced hypocotyl elongation response and the capacity to acclimate to low and high PAR would be a good diagnostic to faithfully classify a plant as shade-tolerant (bad adapted to high PAR exposure, well adapted to live under low PAR and poorly responsive to low R:FR) or shade-avoider (well adapted to high PAR, poor performers under low PAR that elongate when exposed to low R:FR).

Our results also unveiled that activation of low R:FR signaling in shade-avoider species such as *A. thaliana* (At-WT) and shade-tolerant *C. hirsuta* plants with mutations causing low R:FR hypersensitivity (Ch-*sis1* and Ch-*hfr1*) regulated photosynthesis at multiple levels. We confirmed that exposure to W+FR caused a substantial decrease in the levels of photosynthetic pigments (chlorophylls and carotenoids) in these lines (Roig-Villanova *et al*., 2007; Bou-Torrent *et al*., 2015; Molina-Contreras *et al*., 2019; Paulisic *et al*., 2020) and proved that the changes had a direct impact on decreasing phytosynthetic activity (Fig. 5). Low R:FR treatments are known to trigger changes in gene expression within minutes (Kohnen *et al*., 2016). These changes, which are often instrumental for altering rapid growth responses, such as hypocotyl or petiole elongation, are usually mediated by PIFs (Hornitschek *et al*., 2009; Galstyan *et al*., 2011; Cifuentes-Esquivel *et al*., 2013; de Wit *et al*., 2015; Gallemi *et al*., 2017). PIFs were also found to regulate longer-term changes in gene expression such as those affecting photosynthetic genes (Fig. 6). Because loss of PIFQ function in the At-*pifq* mutant resulted in a much attenuated response to W+FR compared to At-WT in terms of photosynthetic gene expression (Fig. 6) but it also prevented photosynthetic pigment and activity loss (Fig. 5), we propose that stabilization of PIFQ proteins following low R:FR exposure triggers a reprogramming of photosynthesis-related gene expression that eventually results in lower pigment levels and reduced photosynthetic activity. Based on the results obtained with other mutants (Fig. 5), we speculate that this signaling network is further influenced by factors such as HFR1 and HY5, which prevent PIF binding to target genes by heterodimerization (Hornitschek *et al*., 2009) or competition for promoter binding sites (Toledo-Ortiz *et al*., 2014), respectively.

Concomitant with the described molecular and physiological changes, we discovered that low R:FR treatment of At-WT seedlings triggered ultrastructural changes in the chloroplast endomembrane systems resembling those occurring after transfer to low PAR (Fig. 7). Furthermore, we demonstrated that pre-treatment with W+FR improved photoacclimation to low PAR of At-WT seedlings but hardly had an effect in low R:FR-hypersensitive mutants of *C. hirsuta*, a shade-tolerant species (Fig. 8). Based on these results, we conclude that the chloroplast ultrastructural changes observed in At-WT plants grown under low R:FR are most likely aimed to acclimate their photosynthetic machinery to perform better under low PAR by, for instance, allowing a more efficient energy transfer. This low R:FR signal-mediated photosynthetic acclimation is likely part of an anticipation mechanism for shade-avoider plants to prepare for the foreseeable reduction in PAR associated with shading. Indeed, low R:FR signals are perceived before actual shading takes place and light becomes limiting, and hence they are considered to act as a warning signal that shading might occur (Martinez-Garcia *et al*., 2010; Casal, 2013). When shade-avoider plants such as *A. thaliana* and most crops (including tomato, cereals, or legumes) grow among taller plants or in a forest understory, they will use the low R:FR signals coming from a closing canopy to elongate (to overgrow its neighbors) but also to readapt its photosynthetic machinery to low PAR before actual shading takes place. By contrast, shade-tolerant plants are adapted to grow under dim light and hence photoacclimation to low PAR is hardly improved even when hypersensitive mutants that show shade-avoider responses in terms of elongation (Fig. 3) and photosynthesis (Fig. 6) are pre-exposed to low R:FR (Fig. 8).

While the observed decrease in photosynthetic pigment and activity levels in shade-avoider plants appears to be part of the anticipation mechanism to an eventual reduction in PAR, a too committed response might be detrimental if light conditions change (e.g., if shading does not occur or shade plants become exposed again to direct sunlight). We have previously shown that a compensation mechanism exist that represses the response to low R:FR when the photosynthetic capacity of chloroplasts is compromised (Ortiz-Alcaide *et al*., 2019). The retrograde (i.e. chloroplast-to-nucleus) pathway that adapts low R:FR perception and signaling to the photosynthetic status of the plant involves the antagonistic factors PIFs and HY5, which also participate in retrograde signaling when underground seedlings are illuminated and start their photomorphogenic (i.e. photosynthetic) development (Ruckle *et al*., 2007; Martin *et al*., 2016; Xu *et al*., 2016; Ortiz-Alcaide *et al*., 2019). The balance of positive and negative regulators together with the chloroplast-mediated control of SAS likely contribute to prevent an excessive response to shade, hence preventing photooxidative damage (resulting from light intensity exceeding the photosynthetic capacity of the plant) and facilitating the return to high R:FR conditions if the low R:FR signal disappears (e.g. if a commitment to the shade-avoidance lifestyle is unnecessary). Together, our work demonstrates that regulation of photosynthetic (chloroplast) performance is both an output and an input of the response of plants to shade. Our results therefore contribute to a better understanding of how plants respond to shade, a knowledge that will contribute to optimally grow crop plants closer together or/and under canopies (e.g., in intercropping settings).

## ACKNOWLEDGEMENTS

We thank M^a^ Rosa Rodríguez (CRAG) for technical support, and George Coupland (MPI for Plant Breeding Research, Cologne, Germany), Rubén Alcazar (Universitat de Barcelona, Spain) and Ignacio Rubio (CRAG) for providing mustard seeds. LM received a predoctoral fellowships from *La Caixa Foundation* (INPhINIT fellowship LCF/BQ/IN18/11660004). WQ is a recipient of a predoctoral Chinese Scholarship Council (CSC) fellowship. Our research is supported by grants from MINECO-FEDER (BIO2017-85316-R, and BIO2017-84041-P) and AGAUR (2017-SGR1211, 2017-SGR710 and Xarba) to JFM-G and MRC. We also acknowledge the support of the MINECO for the “Centro de Excelencia Severo Ochoa 2016-2019” award SEV-2015-0533 and by the CERCA Programme / Generalitat de Catalunya.

## AUTHOR CONTRIBUTIONS

MRC and JFMG conceived the original research plan, directed and coordinated the study. LM and MR-C measured and analyzed photosynthetic parameters and pigment levels; SP, IR-V and WQ performed all the other experiments. All authors analyzed their data and discussed the results. MRC and JFM-G wrote the paper with revisions and contributions or/and comments of all other authors.

